# Neural signatures of disordered multi-talker speech perception in adults with normal hearing

**DOI:** 10.1101/744813

**Authors:** Aravindakshan Parthasarathy, Kenneth E Hancock, Kara Bennett, Victor DeGruttola, Daniel B Polley

## Abstract

In social settings, speech waveforms from nearby speakers mix together in our ear canals. The brain unmixes the attended speech stream from the chorus of background speakers using a combination of fast temporal processing and cognitive active listening mechanisms. Multi-talker speech perception is vulnerable to aging or auditory abuse. We found that ∼10% of adult visitors to our clinic have no measurable hearing loss, yet offer a primary complaint of poor hearing. Multi-talker speech intelligibility in these adults was strongly correlated with neural phase locking to frequency modulation (FM) cues, as determined from ear canal EEG recordings. Combining neural temporal fine structure (TFS) processing with pupil-indexed measures of cognitive listening effort could predict most of the individual variance in speech intelligibility thresholds. These findings identify a confluence of disordered bottom-up and top-down processes that predict poor multi-talker speech perception and could be useful in next-generation tests of hidden hearing disorders.

## Introduction

Slow fluctuations in the sound pressure envelope are sufficient for accurate speech perception in quiet backgrounds (Shannon *et al*., 1995). However, envelope cues are of limited use when speech is embedded in fluctuant backgrounds comprised of multiple talkers, environmental noise or reverberation (Zeng *et al*., 2005). Under these conditions, segregating a target speech stream from background noise requires accurate encoding of low-frequency spectral and binaural cues contained in the stimulus temporal fine structure (sTFS) (Lorenzi *et al*., 2006; Hopkins and Moore, 2009). Monaural sTFS cues convey acoustic signatures of target speaker identity based on the arrangement of peaks in the sound spectrum (e.g., formant frequencies of target speech), while binaural sTFS cues can support spatial separation of target and competing speakers via interaural phase differences (Moore, 2014). With aging and hearing loss, monaural and binaural sTFS cues become less perceptually available, even when audibility thresholds for low-frequency signals that convey sTFS cues are normal (Buss, Hall and Grose, 2004; Fuellgrabe, Moore and Stone, 2015; Leger, Moore and Lorenzi, 2012; Lorenzi *et al*., 2009; Mehraei *et al*., 2014; Strelcyk and Dau, 2009). The biological underpinnings for poor sTFS processing with aging or hearing impairment are unknown, but may reflect the loss of auditory nerve afferent fibers, which degenerate at the rate of approximately 1,000 per decade, such that only half survive by the time a typical adult has reached 40 years of age (Makary *et al*., 2011; Wu *et al*., 2018). A selective loss of cochlear afferent fibers would not be expected to affect audibility thresholds, but could adversely affect the ability of the auditory system to “listen in the dips”, by fully exploiting monaural and binaural sTFS cues available during brief gaps in fluctuant noise levels (Deroche *et al*., 2014; Hopkins, Moore and Stone, 2008; Jin and Nelson, 2010; Moore and Glasberg, 1987; Qin and Oxenham, 2003; Lopez-Poveda and Barrios, 2013).

Excellent processing of a target speaker in a multi-talker background reflects a harmony between high-fidelity encoding of bottom-up acoustic features such as sTFS alongside cognitive signatures of active listening including attention, listening effort, memory, multisensory integration and prediction(Best *et al*., 2009; Narayan *et al*., 2007; Pichora-Fuller *et al*., 2016; Gordon-Salant and Cole, 2016; Winn, Edwards and Litovsky, 2015). These top-down assets can be leveraged to compensate for poorly resolved bottom-up sensory cues, suggesting that listeners with clinically normal hearing that struggle to process speech in noise might be identified by an over-reliance on top-down active listening mechanisms to de-noise a corrupted afferent speech input (Winn, Edwards and Litovsky, 2015; Ohlenforst *et al*., 2017; Besser *et al*., 2015). Here, we apply parallel psychophysical and neurophysiological tests of sTFS processing in combination with physiological measures of effortful listening to converge on a set of neural biomarkers that identify poor multi-talker speech intelligibility in adults with clinically normal hearing.

## Results

### Many individuals seek medical care for poor hearing but have no evidence of hearing loss

We identified the first visit records of English-speaking adult patients from the Mass. Eye and Ear audiology database over a 16-year period, with complete bilateral audiometric records at six octave frequencies from 250 Hz to 8000 Hz according to the inclusion criteria in Figure 1A. Of the 106,787 patient records that met these criteria, we found that approximately one out of every five individuals had no clinical evidence of hearing loss, defined as thresholds > 20 dB HL at test frequencies up to 8 KHz (19,952, 19%, Figure 1B). The majority of these individuals were between 20-50 years old (Figure 1C) and had no conductive hearing impairment, nor focal threshold shifts or “notches” in their audiograms greater than 10 dB (Fig. 1 – Fig. supplement 1A). The thresholds between their left and right ears were also symmetrical within 10 dB for >95% of these patients (Fig. 1 – Fig. supplement 1B). Despite these clinically normal measures of hearing, 45% of these individuals presented to the clinic reporting a primary complaint of decreased hearing or hearing loss (Figure 1D). Absent any objective measure of hearing difficulty, these patients are typically informed that their hearing is “normal” and that they are not expected to experience communication problems.

**Figure 1.**
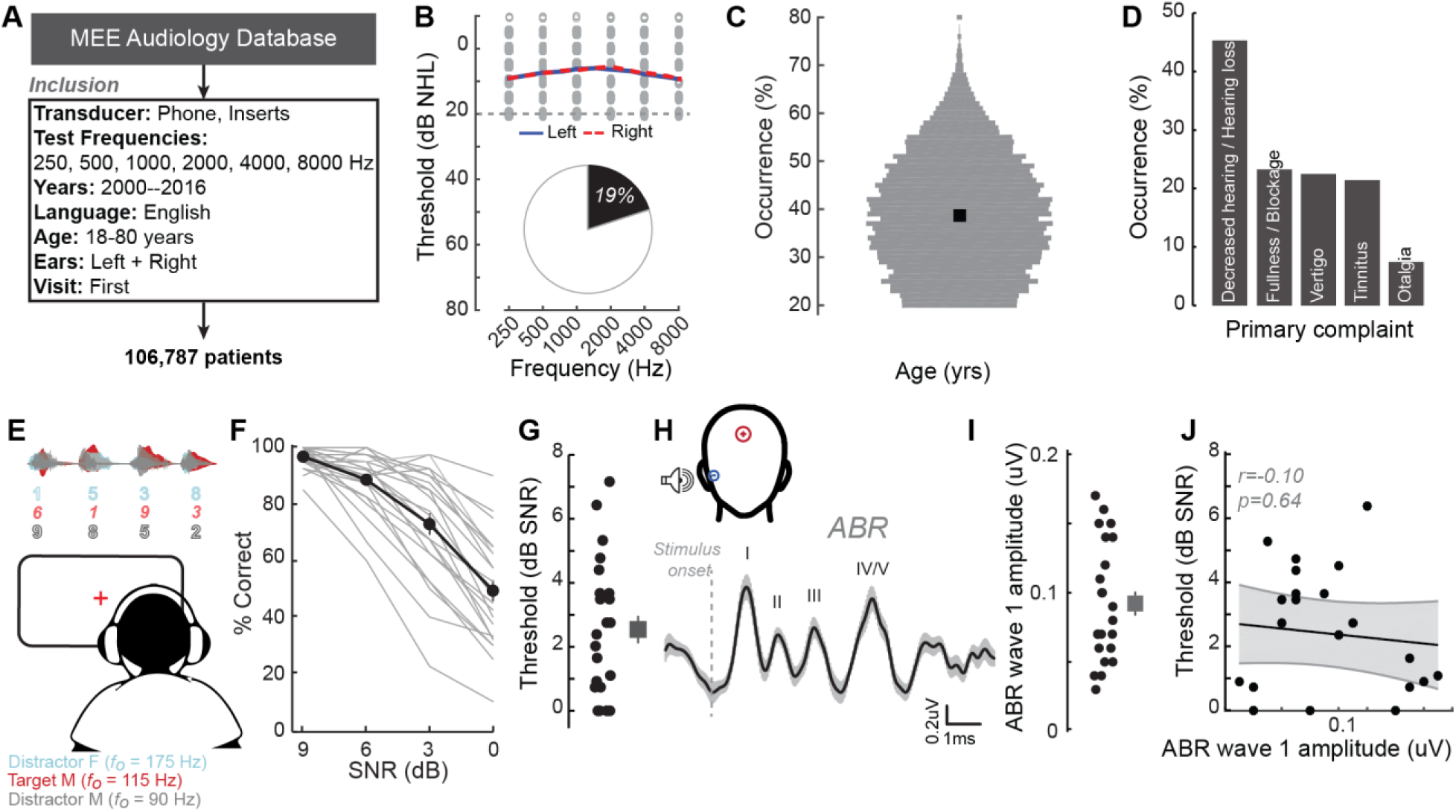
A normal audiogram does not guarantee robust communication in everyday settings. **(A)** Screening criteria for eligible audiology patient records from our hospital collected between 2000 and 2016. **(B)** Bilateral normal audiograms, defined as thresholds better than 20 dB HL (gray dashed line) were identified in 19% of the total patient population. Average audiograms from the left (blue) and right (red) ears are shown with individual data points in gray open circles. **(C)** Normalized age distribution of patients with bilateral normal audiograms shows a larger percentage of younger and middle-aged patients between 20-50 years of age. Black square indicates median age of 39 years. **(D)** Top five primary complaints that resulted in the visit to the clinic for these patients, including perceived hearing loss or decreased hearing presenting in 45% of these patients. **(E)** Schematic of a multi-talker digit recognition task. Subjects (N=23) were familiarized with a target male speaker (red) producing four digits between 1 and 9 (excluding the bi-syllabic ‘7’), while two spatially co-localized distractors, one male and one female, with F0 frequencies above and below the target speaker simultaneously spoke 4 digits at varying signal-to-noise ratios (SNRs). **(F)** Accuracy decreased as a function of SNR at variable rates and to variable degrees. Correct trials required correctly reporting all four digits. **(G)** Variability in individual speech reception thresholds, defined as the SNR that produced a 70.7% success rate. Value at right represents sample mean ± SEM. **(H)** Auditory brainstem responses measured using ear canal tiptrodes yielded robust wave 1 amplitudes, a marker for auditory nerve integrity. Data reflect mean ± SEM. (**I**) Wave 1 values from individual subjects (left) and mean ± SEM of the sample (right). **(J)** No significant associations were observed between the ABR wave 1 amplitudes and speech reception threshold on the multi-talker digit task. r = Pearson’s correlation, and shaded area indicates 95% confidence intervals of the regression line (black) in figures 1-4.

### Speech-in-noise intelligibility varies widely in individuals with clinically normal hearing

Our database analysis suggested that approximately one in ten adults arrived to our clinic seeking care for reduced hearing, only to be told that their hearing was fine. This is not entirely surprising, as most clinical tests are not designed to capture difficulties with “real world” speech communication problems that likely prompted their visit to the clinic. To better understand the nature of their suprathreshold hearing problems, we recruited 23 young or middle-aged listeners (mean age: 29±1 years) that matched the clinically normal hearing from the database profile (Fig. 1 – Fig. supplement2A). Our subjects agreed to participate in a multi-stage research study consisting of self-reported questionnaires, behavioral measures of hearing, and EEG measures of auditory processing (Fig. 1 – Fig. supplement 3).

Although their audiograms were normal and tightly clustered, many of our subjects reported a wide range of difficulties with speech intelligibility, particularly in listening conditions with multiple overlapping speakers (Fig. 1 – Fig. supplement 4A). We directly measured speech-in-noise intelligibility with a digits comprehension task, which simulates the acoustic challenge of focused listening in multi-talker environments, while eliminating linguistic and contextual speech cues. Subjects attended to a familiar male speaker (F0 = 115 Hz) producing a stream of four digits in the presence of one male and one female distracting speakers (F0 = 90Hz and 175 Hz, respectively). The distracting speakers simultaneously produced digits at variable signal-to-noise ratios (SNRs) (Figure 1E). Performance on the digits task was highly variable, especially at SNRs of 3 and 0 dB (Figure 1F). Speech reception thresholds, defined as the 70.7% correct point on the response curve, varied widely across a 0-7 dB SNR range (Figure 1G). We found that speech intelligibility thresholds were significantly correlated with the subjects’ self-reported difficulties in multi-speaker conditions, suggesting that the digits comprehension task captures aspects of their real-world communication difficulties (r = 0.46, p = 0.02, Fig. 1 – Fig. supplement 4B).

### Peripheral markers of noise damage do not explain performance on the speech-in-noise task

We first determined whether simple adaptations of existing clinical tests could identify deficits in multi-talker speech segregation. We measured hearing thresholds at extended high frequencies, a marker for early noise damage (Fausti *et al*., 1981; Mehrparvar *et al*., 2011; Le Prell *et al*., 2013). Despite their clinically normal audibility at lower frequencies (250-8000Hz), subjects exhibited substantial variability in their extended high frequency thresholds (Fig. 1 – Fig. supplement 2B). We also noted considerable inter-subject variability in the amplitude of auditory brainstem response (ABR) wave 1, a marker for peripheral nerve health, which decreases over the course of normal aging or following neuropathic noise exposure (Figure 1H-I) (Parthasarathy and Kujawa, 2018; Fernandez *et al*., 2015). Both indirect measures of auditory peripheral health showed substantial inter-subject variability despite normal hearing thresholds, but the particular levels were not statistically related to performance on the competing digits task, demonstrating that these markers of peripheral damage do not have a direct contribution to this measure of multi-talker speech performance in our subject population (r = 0.10 p = 0.64, Figure 1J, Fig. 1 – Fig. supplement 2D).

### Encoding of sTFS cues predicts speech-in-noise intelligibility

Poor processing of sTFS cues has long been associated with elevated speech recognition thresholds, especially in patients with hearing loss (Lorenzi *et al*., 2006). In a classic test of sTFS processing, subjects are asked to detect a slow, subtle FM imposed on a low-frequency carrier (Moore and Skrodzka, 2002; Moore and Sek, 1996; Moore and Sek, 1995; Sek and Moore, 1995). We tested our subjects with this psychophysical task, which uses an adaptive two-interval forced choice procedure to converge on the threshold for detecting FM of a 500 Hz tone (Figure 2A). FM detection thresholds varied widely between subjects (Figure 2B) and were strongly correlated with performance on the competing digits task (r = 0.85, p<0.001, Figure 2C). We were struck that detection thresholds for such a simple stimulus could accurately predict performance in a much more complex task. On the one hand, sensitivity to FM could reflect a superior low-level encoding of sTFS cues that are critical for segregating a target speech stream from distractors. Alternatively, perceptual thresholds for FM could reflect a superior abstracted representation of the stimulus features at any downstream stage of neural processing, and not the high fidelity representation of sTFS cues, per se. Taking this line of argument a step further, a correlation between competing talker thresholds and FM thresholds may not reflect the stimulus representation at all, but instead could reflect subjects’ general aptitude for utilizing cognitive resources such as attention and effort to perform a wide range of listening tasks (Pichora-Fuller *et al*., 2016).

**Figure 2.**
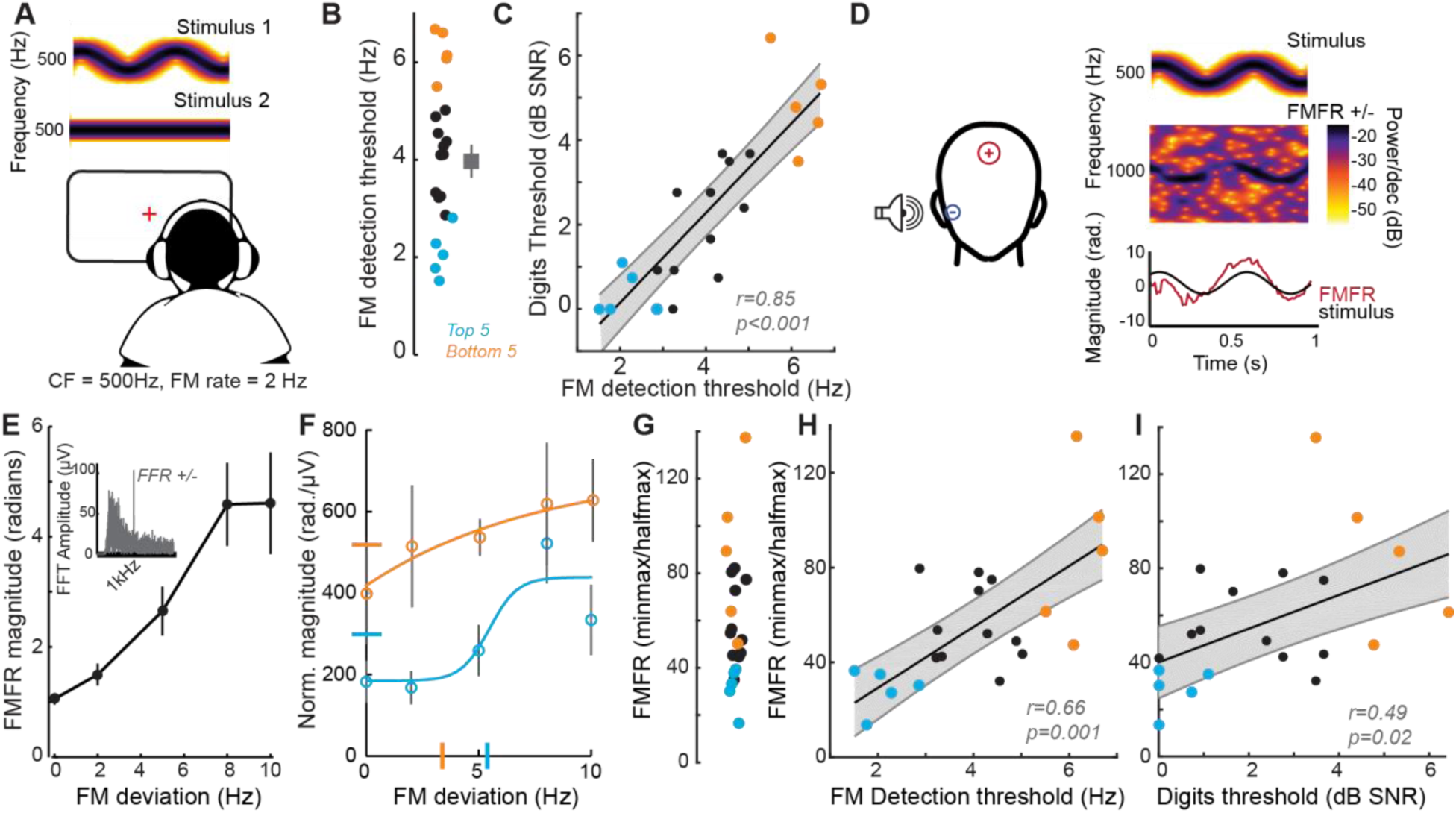
Perceptual and neural processing of sTFS cues predict speech in noise intelligibility. **(A)** Design of a psychophysical task to measure frequency modulation (FM) detection threshold. Participants indicated FM in a 500Hz tone in an adaptive (2-down 1-up) 2 alternative forced choice task. **(B)** Individual (left) and average ± SEM (right) FM detection thresholds. Top and bottom 5 performers (∼20^th^ percentile) on the behavioral FM detection task are shown in blue and orange respectively, in panels B-C and F-I **(C)** FM detection thresholds were strongly predictive of speech in noise recognition threshold defined with the multi-talker digit comprehension task. **(D)** An objective neurophysiological measure of monaural sTFS processing was obtained using ear canal (-) and scalp (+) electrodes, and a 500Hz tone presented with various FM deviations in alternating polarity. The averaged response was analyzed at 1000Hz (2F) in order to minimize contributions by the cochlear microphonic and emphasize neural generators. The magnitude of the FM following response (FMFR) was computed using a heterodyne. **(E)** The FMFR magnitude increased as a function of FM deviation up to ∼8 Hz. *Inset:* The FMFR magnitude was normalized by the pure tone phase-locking amplitude of each subject to minimize variability due to head size and recording conditions. **(F-G)** A sigmoidal fit to the normalized FMFR growth curve was used to calculate an FMFR measure of slope for each subject, by dividing the overall dynamic range of the response by the halfway point to the maximum (illustrated in (*F)* for the top and bottom 5 performers of the behavioral task). Blue and orange bars indicate the X and Y axes intercepts of the halfway point of the fit. **(H-I)** The neurophysiological FMFR was strongly predictive of FM detection thresholds determined with the psychophysical task (*H*) as well as performance on the digits comprehension task (*I*).

We developed a direct physiological measure of early neural processing of FM cues to test the hypothesis that low-level encoding of acoustic sTFS cues is associated with superior speech-in-noise processing. We determined that an ear canal to Fz electrode montage was sensitive to evoked potentials generated by the auditory nerve (Fig. 1I, Fig. 2 – Fig. supplement 1A-B), but we also wanted to exclude any pre-neural contributions, such as the cochlear microphonic, which is generated by outer hair cells (Fettiplace, 2017). Since neural responses at the auditory nerve act as a half-wave rectifier for sinusoidal inputs, we used FM tones of alternating polarity and analyzed responses at twice the carrier frequency. This approach allowed us to minimize non-rectifying pre-neural components and emphasize neural phase-locking (Lichtenhan *et al*., 2014). We observed robust phase-locked following response to the FM stimulus (Figure 2D, termed the FM following response or FMFR). We used a heterodyne method to extract the FMFR for FM depths up to 10 Hz (Figure 2E). To factor out the variability due to head size and overall electrode SNR we calculated the amplitude of the carrier frequency following response to a tone with 0 Hz FM and then expressed the FMFR magnitude as a fraction of this value (Figure 2E, **inset**). Sigmoidal fits to the FMFR growth function (illustrated for the top and bottom 5 performers on the behavioral task in Figure 2F) were further reduced to a single value per subject by dividing the maximum dynamic range for each subject (min-max) by the halfway point to get a measure of slope (halfmax; Figure 2G). With this approach, subjects with a wide dynamic range for encoding FM depth have more robust FM encoding and therefore smaller min-max/halfmax ratio values.

We found that robust low-level encoding of FM cues was highly predictive of an individual’s performance on the FM psychophysical detection task (r = 0.66 p = 0.001; Figure 2H), suggesting that the FMFR can provide an objective neurophysiological measure of an individual’s ability to encode sTFS cues. Importantly, FMFR also predicted performance on the competing digits task (r = 0.49, p = 0.02, Figure 2I).

These data suggest that the strong association between psychophysical tests for FM detection and speech-in-noise intelligibility can be attributed, at least in part, to encoding of FM cues at the earliest stages of auditory processing.

### Encoding of unrelated sTFS cues do not predict speech-in-noise intelligibility

We reasoned that the correlation between low-level FM encoding and speech intelligibility might just reflect a correlation between any measure of fast temporal processing fidelity and speech intelligibility. This could be addressed by measuring temporal processing fidelity on an unrelated stimulus and noting whether it had any correlation with speech-in-noise thresholds. Interaural timing cues can improve speech processing in noise, but would not be expected to have any association with the competing digits task used here, where the identical waveform was presented to both ears. To test whether poor encoding of binaural sTFS cues would also predict poor performance in the competing digits task, we performed parallel psychophysical and electrophysiological measurements of sensitivity to interaural phase differences (McAlpine *et al*., 2016; Undurraga *et al*., 2016; Haywood *et al*., 2015; Ross *et al*., 2007).

In this task, the phase of a 500Hz tone presented to each ear was shifted by a variable amount (up to 180°), creating the percept of a tone that moved from the center to the sides of the head. To eliminate phase transition artifacts, an amplitude modulation of ∼41 Hz was imposed on the tone, such that the instantaneous phase shift always coincided with the null of the amplitude envelope (Figure 3A) (Undurraga *et al*., 2016; Haywood *et al*., 2015). To test psychophysical thresholds for binaural sTFS cues, subjects indicated the presence of an interaural phase difference (IPD) in one of two tokens. IPD thresholds were variable, ranging between 5 and 25 degrees (Figure 3B). To quantify electrophysiological encoding of IPD, recordings were made with electrodes in a vertical Fz-C7 montage to emphasize binaural generators (Fig. 2– Fig. supplement 1B). The IPD was alternated at 6.8 Hz, inducing a following response to the IPD (IPDFR) as well as a phase-insensitive envelope following response at the amplitude modulation rate (Figure 3C). As expected, the amplitude of the IPDFR at 6.8 Hz increases with larger interaural time differences, whereas the amplitude of the 40 Hz envelope following response remains constant (Figure 3D). As above, we minimized variability due to head size and electrode SNR by expressing the IPDFR amplitude as a fraction of the envelope following response. Sigmoidal fits to the normalized growth curves were then used to calculate min-max/halfmax values, similar to the FMFRs (shown for the top and bottom 5 performers on the behavioral task in Figure 3E). Like the FMFR above, we noted a strong association between an individual’s psychophysical threshold for IPD and the growth of the electrophysiological IPDFR (r = −0.65 p = 0.001, Figure 3F). Unlike the FMFR, subjects that were most sensitive to IPD showed a large, rapid increase in IPDFR amplitude across the testing range resulting in a large min-max and a small half-max (Figure 3E). As a result, the correlation between psychophysical threshold and IPDFR is negative (Figure 3F) whereas the correlation between FM threshold and the FMFR amplitude is positive (Figure 2H). This can be attributed to inherent differences in the FMFR (a measure of sTFS phase locking) versus the IPDFR (a measure of sTFS envelope processing; see Discussion). More to the point, neither the psychophysical threshold, nor the IPDFR amplitude had statistically significant correlations with the digits in noise threshold, confirming that task performance was specifically linked to encoding of task-relevant FM cues and not a general sensitivity to unrelated sTFS cue (IPD threshold and speech, r = 0.21 p = 0.34; IPDFR and speech, r = 0.20 p = 0.38, Figure 2G).

**Figure 3.**
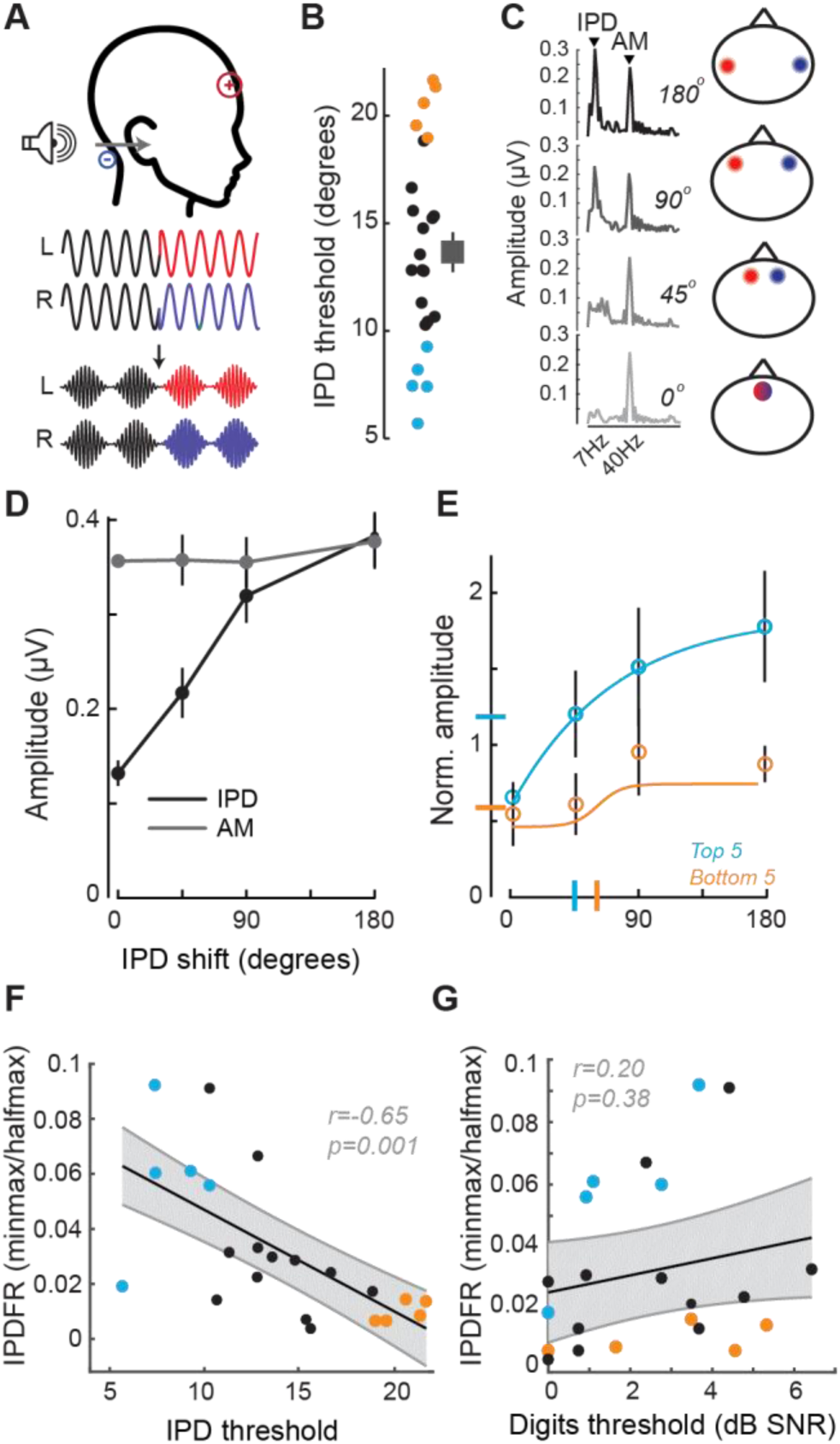
Neural and perceptual processing of rapid temporal cues unrelated to the speech task do not predict speech recognition thresholds. **(A)** Design of interaural phase difference (IPD) detection task. The phase of a 500Hz tone instantaneously shifted from diotic (aligned interaural phase) to dichotic (variable interaural phase). Amplitude modulation (AM) at 40.8 Hz was aligned to the interaural phase shift such that the amplitude minimum coincided with the phase transition. **(B)** Minimal IPD detection threshold was measured in a 2 alternative forced choice task. IPD thresholds varied between 5 and 25 degrees across individual subjects (left), mean ± SEM shown at right. Top and bottom 5 performers (∼20^th^ percentile) on the behavioral FM detection task are shown in blue and orange respectively. **(C-D)** In EEG recordings, the IPD alternated between diotic and dichotic at a rate of 6.8 Hz. Fast Fourier transforms of scalp-recorded evoked responses revealed a phase-dependent IPD following response (IPDFR) at 6.8 Hz and a 40.8 Hz AM envelope following response. **(E)** The IPDFR magnitude was expressed as a fraction of the envelope following response for each subject to minimize variability due to head size and recording conditions. Min-max and half-max values were computed from sigmoidal fits to the normalized IPDFR growth function (Illustrated here for the top and bottom 5 performers on the behavioral task) **(F-G)** The IPDFR was strongly predictive of IPD detection thresholds (*F*), but not performance on the digits comprehension task (*G*).

### Pupil-indexed effortful listening predicts speech intelligibility

Speech recognition is a whole brain phenomenon that is intimately linked to cortical processing as well as cognitive resource allocation such as listening effort, spatial attention, working memory, and prediction (Shinn-Cunningham, Best and Lee, 2017; Ding and Simon, 2012; O’Sullivan *et al*., 2015; Ruggles, Bharadwaj and Shinn-Cunningham, 2011; Mesgarani and Chang, 2012). In this sense, encoding of bottom-up cues such as sTFS can provide critical building blocks for downstream speech processing but ultimately provide an incomplete basis for predicting performance on cognitively demanding listening tasks. To capture variability in speech processing that was not accounted for by sTFS cues, we measured task-evoked changes in pupil diameter, while subjects performed the digits comprehension task. Under isoluminous conditions, pupil diameter can provide an objective index of the sensory and cognitive challenge of processing a target speech stream in the presence of distracting speakers (Zekveld, Kramer and Festen, 2010; Wang *et al*., 2018; Wang *et al*., 2017; Winn, 2016; Winn, Edwards and Litovsky, 2015; Koelewijn *et al*., 2015). Prior work has shown that increased pupil dilation in low SNR listening conditions can reflect greater utilization of top-down cognitive resources to enhance attended targets, whereas smaller pupil changes have been associated with higher fidelity bottom-up inputs that do not demand additional listening effort to process accurately (Zekveld *et al*., 2014; Zekveld and Kramer, 2014; Koelewijn *et al*., 2012) (Figure 4A). We confirmed here that the fractional change in pupil diameter was linearly related to the SNR of the target speaker (Figure 4B) in 16 subjects that provided measurable pupil signals (see Methods for a statement on exclusion criteria). In the same spirit as removing variability related to head size and electrode SNR, we factored out unrelated measurement noise by expressing the SNR-dependent change in pupil diameter as a fraction of the light-induced pupil change in each subject (Figure 4C). We confirmed that individuals with steeper pupil recruitment functions had more difficulty in the multi-talker speech task, leading to a significant correlation between half-max/min-max pupil change and speech intelligibility threshold (r = 0.53, p = 0.03, Figure 4D).

**Figure 4.**
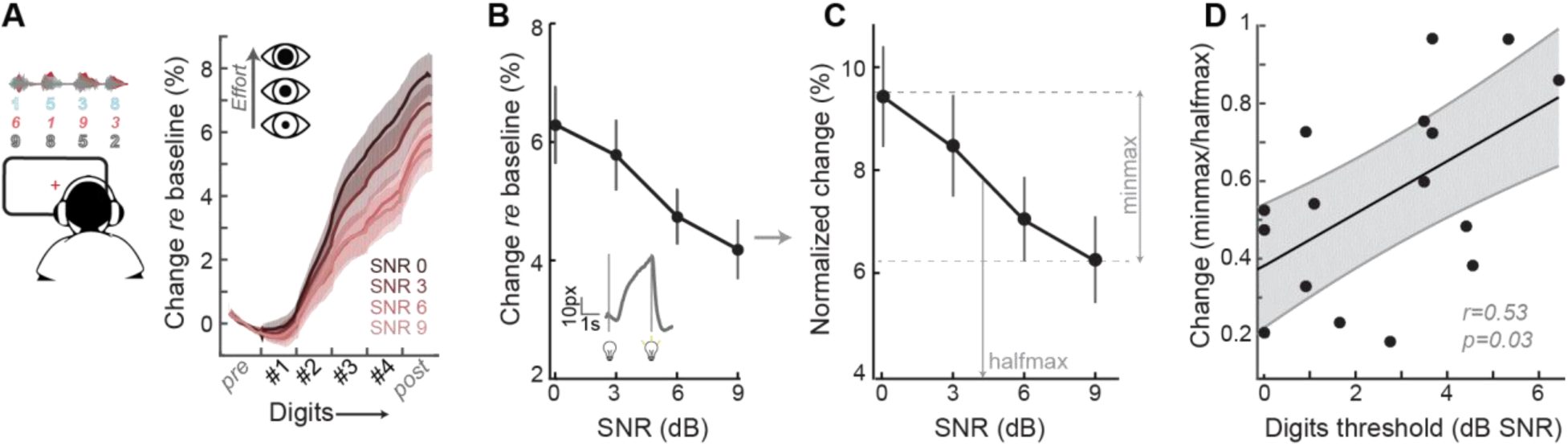
A pupil-indexed measure of effortful listening predicts multi-talker speech recognition thresholds. **(A)** Fractional change in pupil diameter was measured under isoluminous conditions before, during and after the 4-digit sequence at various SNRs. **(B)** The peak fractional change in pupil diameter was normalized to the light-induced pupil change for each SNR **(C).** The SNR-dependent change in pupil diameter was calculated as the min-max/halfmax. **(D)** Greater recruitment of pupil-indexed effortful listening across SNRs was significantly associated with the speech intelligibility threshold. Baseline changes in pupil across the testing session, taken as a measure of listening fatigue, showed no relationship with task performance (Fig. 4 – Fig. supplement 1).

### Predictors of speech intelligibility – shared and private variability

Our overall motivation was to develop objective physiological markers that might explain complaints of poor speech communication in individuals with clinically normal hearing. Here, we examined whether poor speech-in-noise intelligibility was associated with poor auditory nerve integrity (indexed here by ABR wave 1 amplitude), poor encoding of monaural sTFS cues (as indexed by the FMFR), generally poor fast temporal processing (indexed here by IPDFR) and increased utilization of cognitive resources related to effortful listening (indexed here by pupil change). Importantly, none of these indices were correlated with each other, indicating that - in principle – each of these markers could account for statistically independent components of the total variance in speech performance (Figure 5A**, right**). In practice, only FMFR and pupil showed a significant independent correlation with speech intelligibility threshold (Figure 5A**, left**).

**Figure 5.**
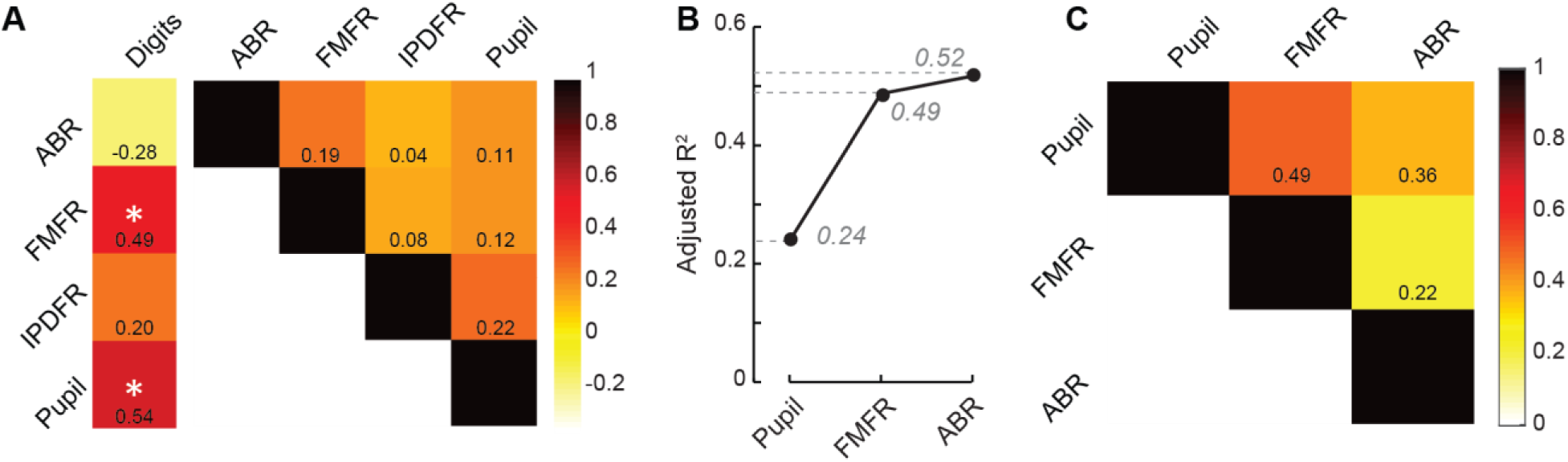
A multiple variable linear model of bottom-up and top-down neural markers best predicts speech intelligibility thresholds. **(A)** The four neural markers studied here (ABR wave 1, FMFR, IPDFR and pupil-indexed effortful listening) were not correlated with each other. FMFR and Pupil were both significantly correlated with the outcome measure, digits comprehension threshold. White asterisk indicates p < 0.05 with a univariate linear regression model. **(B)** A multivariate regression model measuring the adjusted R2 (proportion of variance explained by predictors, adjusted for the number of predictors in the model) reveals incremental improvement in the prediction of the digits comprehension threshold when pupil, FMFR and ABR wave 1 amplitudes are added in succession. Adding additional neural markers to the model did not improve the total explained variance. **(C)** All two-variable combinations were analyzed to study order effects for introducing variables into the model. The combination of pupil diameter and FMFR was still the most optimal model for explaining variance on the speech-in-noise task. Numbers indicate adjusted R2 values for each combination.

To determine whether combining these independent metrics could account for an even greater fraction of the total variance, we used a multiple variable linear regression model, and computed the adjusted R^2^ values, after adding each successive variable. Variables were added in decreasing order of individual R^2^ values. The adjusted R^2^ penalizes for model complexity incurred due to the addition of more variables (See methods). With this model, listening effort indexed by pupil diameter explained 24% of the variance in the digit comprehension task. Adding in monaural fine structure processing measured using the FMFR increased the adjusted R^2^, explaining 49% of the overall variance. Adding in the ABR wave 1 only provided a marginal increase in predictive power, raising the total explained variance to 52% (Figure 5B). Adding additional neural markers such as the IPDFR or extended high frequency thresholds did not provide any further increase in the overall variance explained. Among the neural measures studied here, the best linear model for speech intelligibility included a measure of bottom-up monaural fine structure processing and a measure of top-down listening effort. In order to account for order effects in the model, we also looked at the adjusted R^2^ for all 2-variable combinations between the FMFR, pupil diameter and the ABR. The combination of FMFR and pupil diameter provided the best model in all order configurations (Figure 5C). Finally, even though the behavioral FM detection thresholds and the FMFR were correlated (Figure 2H), constructing a model with both the behavioral and physiological measure, along with pupil diameter increased the variance explained to 78%, suggesting that the behavioral FM detection task reflects an additional aspects of auditory processing that is not captured by the combination of peripheral sTFS encoding and non-sensory measures of listening effort.

## Discussion

### Neural and perceptual processing of temporal fine structure

The cochlea acts as a limited-resolution filter bank that breaks down the broadband speech waveform into a spatially organized array of narrowband signals. Each cochlear filter contains two types of information that are encoded and reconfigured by neurons within the spiral ganglion and central auditory pathway: sTFS and stimulus temporal envelope. STFS cues consist of rapid oscillations near the center of each cochlear filter that are encoded by phase-locked auditory nerve action potential timing (Henry and Heinz, 2013). Envelope cues, by comparison, reflect slower changes in amplitude over time that can be encoded by the short-term firing rate statistics of auditory nerve fibers (Joris and Yin, 1992). The ability to detect slow rates of FM (<∼6Hz) at low carrier frequencies (<∼1500Hz) has long been associated with sTFS processing (Moore and Sek, 1995; Sek and Moore, 1995). Under these stimulus conditions, changes in FM deviations are hypothesized to be conveyed by spike timing information within a cochlear filter (Moore and Sek, 1995).

The strong correlation between detection threshold for pure tone FM and speech recognition threshold is striking and has now been documented by several independent groups using a variety of speech-on-speech masking paradigms, but not with non-speech maskers (Whitton *et al*., 2017; Strelcyk and Dau, 2009; Johannesen *et al*., 2016). The exact mechanism of FM coding by the auditory pathway is not entirely clear, with some studies suggesting that these FM cues are converted to amplitude modulation cues in the early stages of auditory processing, and hence that the perception of FM relies more on neural envelope cues rather than the neural phase locking to sTFS (Ghitza, 2001; Whiteford, Kreft and Oxenham, 2017; Whiteford and Oxenham, 2015). Here, we characterized processing of FM tones using a combination of classic psychophysical tests and newly developed ear canal EEG following responses. Because we looked at changes in phase-locking to the carrier, and not the actual rate of FM, we were able to explicitly emphasize neural coding of these timing cues in the early auditory system, while minimizing contributions from the recovered envelope, which would be reflected as the 2Hz FM rate. The FMFR was correlated with the behavioral FM detection, suggesting that this response reflects aspects of the behavioral FM detection task (Figure 2H), and was also correlated with performance on the digits task (Figure 2I), suggesting that the representation of these fine stimulus timing cues contributes to speech-in-noise intelligibility.

Subjects with the lowest FM detection thresholds exhibited a small increase in FMFR amplitudes across a broad range of shallow excursion depths before suddenly increasing at FM excursion depths that exceeded the limits of a single cochlear filter, perhaps indicating the transition from a timing to a place code (Figure 2F). By contrast, the IPDFR transfer function in subjects with the lowest IPD thresholds increased steadily for all IPDs above zero (Figure 3D). As a result, top psychophysical performers had a shallow transfer function for FM excursion depth but a steep transfer function for IPDFR, producing a positive correlation between FMFR and FM detection threshold (Figure 2H) and a negative correlation between the IPDFR and IPD detection threshold (Figure 3F). As described above, the FMFR is calculated as the phase coherence to the FM carrier, whereas the IPDFR is calculated as the entrainment to the rate of IPD alternation. As these measures are in no way equivalent, there is no reason to expect the same relationship between each transfer function and psychophysical detection limits.

### From mechanisms to biomarkers for hidden hearing disorder

Patients with bilateral normal audiograms represented ∼19% of the patient population at the Mass. Eye and Ear Infirmary, 45% of whom reported some form of perceived hearing loss as their primary complaint (Figure 1D). The combination of an increased lifespan and the increased use of in-ear listening devices will likely exacerbate the societal impact of this sensory disorder, leaving hundreds of millions of people straining to follow conversations in noisy, reverberant environments typically encountered in the workplace and social settings (Goman and Lin, 2016; Lin *et al*., 2011; Hind *et al*., 2011; Ruggles, Bharadwaj and Shinn-Cunningham, 2012; Ruggles, Bharadwaj and Shinn-Cunningham, 2011). Hearing aids and other in-ear amplification devices are of limited use for persons with clinically normal hearing thresholds who primarily complain of difficulty listening in situations involving noise and competing messages (Bentler and Duve, 2000).

From a health care perspective, the issue is more than just a lack of treatment options, as conventional clinical tests for hearing health do not address processing of suprathreshold communication signals in noise, but instead are primarily focused on identifying minimally audible levels of pure tones or isolated words in silence. Low-channel EEG systems already exist in most hearing health clinics for ABR measurements, which could be adapted to the FMFR test described here. Pupillometry systems are also relatively low cost and straightforward to use, suggesting that the combination of FMFR and pupil-indexed listening could theoretically serve as clinically useful biomarkers for disordered speech-in-noise processing. Identifying non-invasive physiological correlates of poor speech intelligibility in noise would be valuable in the development of next-generation hearing health measures but might also shed light on the modes of biological failures underlying this common sensory disorder.

The mechanism for poor encoding of sTFS cues remains to be determined, though one possibility is that it reflects a loss of cochlear afferent neurons that synapse onto inner hair cells. Auditory nerve fiber loss has been observed in cochlear regions with normal thresholds in many animal species as well as post-mortem analysis of human temporal bone specimens (Wu *et al*., 2018; Valero *et al*., 2017; Viana *et al*., 2015; Furman, Kujawa and Liberman, 2013; Kujawa and Liberman, 2009). In humans, recent findings suggest that appreciable auditory nerve fiber loss begins in early adulthood, well before degeneration is noted in cochlear sensory cells or spiral ganglion cell bodies (Wu *et al*., 2018). In animal models, a loss of cochlear afferent synapses disrupts the encoding of rapid timing cues, without affecting thresholds, consistent with observations made in our subject cohort (Parthasarathy and Kujawa, 2018; Shaheen, Valero and Liberman, 2015). In humans, it is impossible to directly assess the status of cochlear afferent synapses in vivo, though indirect proxies for cochlear afferent innervation may be possible (Liberman *et al*., 2016; Mehraei *et al*., 2016; Bharadwaj *et al*., 2015; Guest *et al*., 2017; Prendergast *et al*., 2017a; Prendergast *et al*., 2017b; Grinn *et al*., 2017; Bramhall *et al*., 2017). Prior work has emphasized the amplitude of ABR wave 1 and extended high frequency hearing thresholds as possible indirect markers of cochlear synapse loss. We observed considerable individual variability in both of these measures, although this variance had no statistical relationship with speech-in-noise thresholds in our subjects (Figure 1J, Figure1-Figure supplement 2D).

Speech perception does not arise directly from the auditory nerve, but rather reflects the patterning of neural activity in the central auditory pathway. Therefore, one might expect a correlation between a well-designed proxy for auditory nerve integrity and a behavioral measure of speech recognition accuracy, but the correlation would never be expected to be too high simply because the periphery is a distal – not proximal – basis for speech perception. Hearing loss profoundly affects gene expression, cellular morphology, neurotransmitter levels and physiological signal processing at every stage of the central pathway - from cochlear nucleus to cortex - and these central sequelae resulting from a peripheral insult would also be expected to affect the neural representation of speech in ways that cannot be accounted for purely by peripheral measures (Caspary *et al*., 2008; Chambers *et al*., 2016; Sarro *et al*., 2008; Parthasarathy, Herrmann and Bartlett, 2019; Auerbach, Radziwon and Salvi, 2019). To this point, adding a non-peripheral measure of pupil-indexed listening effort improved the correlation with speech recognition thresholds. Moreover, the correlation between the psychophysical FM detection thresholds was more highly correlated with speech recognition than the neural measure of low-level FM encoding, suggesting that the behavioral task captured additional aspects of FM detection not present in the FMFR. We would expect that an additional neural marker of higher-order stream segregation (Lu *et al*., 2017; Shamma, Elhilali and Micheyl, 2011; Teki *et al*., 2013; Krishnan, Shamma and Elhilali, 2014) or direct neural speech decoding (Mesgarani *et al*., 2014; Pasley *et al*., 2012; Mesgarani *et al*., 2008; Presacco, Simon and Anderson, 2016; Ding and Simon, 2012; Ding and Simon, 2009) would complement our measures of cognitive active listening (via pupil) and low-level sTFS processing (via FMFR). Future studies could explore how additional neural markers of downstream central processing could be combined with the markers for auditory cognition and low-level sTFS processing described here to converge on an even better aggregate model for predicting individual variability in speech-in-noise intelligibility.

## Methods

### Subjects

All procedures were approved by the institutional review board at the Massachusetts Eye and Ear Infirmary. Twenty seven subjects (13 male, 14 female) were recruited and provided informed consent to be tested as part of the study. Of these, 4 subjects were excluded for either failing to meet the inclusion criteria (1 male, 1 female, see below for inclusion criteria) or not completing more than 60% of the test battery (2 male). One subject (female) did not participate in the electrophysiology tests, but data from the other two sessions were included for relevant analyses. Subjects were compensated per hour for their participation in the study.

### Testing paradigm - Overview

Eligibility of the participants was determined on day of first visit by screening for cognitive skills (Montreal Cognitive Assessment, MOCA >25 for inclusion), depression (Beck’s depression Inventory, BDI <21 for inclusion), tinnitus (Tinnitus reaction questionnaire, TRQ <72 for inclusion), use of assistive listening devices (“Do you routinely use any of the following devices – cochlear implants, hearing aids, bone-anchored hearing aids or FM assistive listening devices” - subjects were excluded if they answered yes to any of the above) and familiarity with English (“Are you a native speakers of English”, and “If not, Are you fluent or functionally fluent in English?” - subjects were excluded if they answered no for both questions). Eligible participants were then tested in a double walled acoustically isolated chamber with an audiometer (Interacoustics AC40, Headphones: TDH39) to confirm normal audiograms with thresholds ≤ 20dB HL for frequencies up to 8 kHz. Participants then performed high frequency audiometry (Headphones: Sennheiser HDA200) and the digits comprehension task paired with pupillometry (described in detail below). Subjects were then sent home with tablet computers (Microsoft Surface Pro 2) and calibrated headphones (Bose AE2). Subjects were asked to complete additional suprathreshold testing (FM detection, IPD detection) and questionnaires - Noise exposure questionnaire (NEQ) (Johnson *et al.,* 2017) Speech, spatial and Qualities of hearing scale SSQ (Gatehouse and Noble, 2004),Tinnitus handicap questionnaires (Newman, Jacobson and Spitzer, 1996) in a quiet environment over the course of 8 days. The microphone on the tablet was used to measure ambient noise level throughout home-based testing. If noise levels exceeded 60 dB A, the participant was locked out of the software, provided with a warning about excessive noise levels in the test environment, and prompted to find a quieter location for testing. Subjects returned to the laboratory on Day 10 (± 1 day) for electrophysiological testing.

### Speech Intelligibility Threshold

Subjects were introduced to the target male speaker (F0 = 115Hz) as he produced a string of four randomly selected digits (digits 1-9, excluding the bisyllabic ‘7’) with 0.68s between the onset of each digit. Once familiarized, the task required subjects to attend to the target speech steam in the presence of two additional speakers (male, F0 = 90Hz; female, F0 – 175 Hz) that produced randomly selected digits with matched target-onset times. The two competing speakers could not produce the same digit as the target speaker or each other, but otherwise digits were selected at random. The target speaker was presented at 65 dB SPL. The signal-to-noise ratio of the distractors ranged from 0-20 dB SNR. Subjects reported the target 4-digit sequence using a virtual keypad on the tablet screen 1s following the presentation of the 4^th^ digit. Subjects were initially provided with visual feedback on the accuracy of their report in four practice blocks comprised of 5 trials each and 4 SNRs (target only, 20, 9 and 3 dB SNR). Testing consisted of 40 blocks of 8 trials each, with SNRs of 9, 6, 3 and 0 dB presented in a randomized order for each cycle of four blocks. The first three trials of each block served as refreshers to familiarize the subject with the target speaker at 20 dB SNR before progressively decreasing to the test SNR presented in the last five trials of each block. Trials were scored as correct if all four digits entered into the keypad matched the target speaker sequence.

### Frequency Modulation Detection Threshold

Subjects were introduced to the percept corresponding to frequency modulation (FM) through a virtual slider on the tablet computer that they manipulated to increase and decrease the FM excursion depth of a 500 Hz tone. High excursions were labeled “squiggly” to allow the subjects to associate the sound with a label that could be used when completing the 2-interval 2-alternative forced choice detection task. After initial familiarization, two tones (carrier frequency = 500 Hz, duration = 1 s, level = 55 dB SL) were presented to subjects with an interstimulus interval of 0.5 s. Frequency modulation was applied at a rate of 2 Hz to one of the two tones (order selected at random) and the other tone had no FM. A quasi-sinusoidal amplitude modulation (6 dB depth) was applied to both tones to reduce cochlear excitation pattern cues (Moore and Sek, 1996). The subject reported whether the first or second tone was “squiggly” (i.e., was the FM tone). A two-down one-up procedure converged on the FM excursion depth that subjects could identify with 70.7% accuracy (Levitt, 1971). FM excursion depth was initially set to 75 Hz and was then changed by a factor of 1.5 for the first 5 reversals, decreasing to a factor of 1.2 for the last 7 reversals. The geometric mean of the last 6 reversals was used to compute the run value. A minimum of 3 runs were collected. The coefficient of variation (standard deviation / mean) for the reversal values was computed during testing. If the coefficient of variation was > 0.2, additional runs were collected until this criterion was met or six runs had been collected, whichever came first. The median threshold value obtained across individual runs defined the participant’s FM detection threshold.

### Interaural Phase Difference Detection Threshold

Sensitivity to interaural phase difference was tested using a 2-interval 2-alternative forced choice task. Sound tokens consisted of tones presented simultaneously to both ears at the same carrier frequency (520 Hz), amplitude modulation rate (100% depth at 40.8 Hz), duration (1s) and level (85 dB SPL, 50 ms raised cosine onset/offset ramps). Each token was separated by a 0.5s silent interval. Both tokens started in phase. But for one of the two tokens, a phase shift was applied to the tone in each ear in opposing polarity, 0.5s after tone onset. This produced a perceptual switch, where the sound “moved” from a diotic to a dichotic percept. The subjects were asked which of two sound tokens “moved” in the middle. Subjects were familiarized with the task in two practice blocks of ten trials each and provided visual feedback about their accuracy in identifying the tone that “moved”. A two-down-one-up procedure was used to converge on the phase shift that could be identified with 70.7% correct accuracy. The phase shift was initially set to 81 degrees and changed by a factor of 1.5 for the first 4 reversals, decreasing to a factor of 1.2 for the last 6 reversals. The geometric mean of the last 6 reversals was used to compute the run value. The criteria for determining the number of runs and the threshold matched the FM detection task above.

### Electrophysiology

EEG recordings were performed in an electrically shielded sound attenuating chamber. Subjects reclined in a chair and were instructed to minimize movements. Arousal state was monitored but not regulated. Most subjects reported sleeping through the recordings. The recording session lasted ∼3 hours and subjects were given breaks as necessary. Recordings were done on a 16-channel EEG system (Processor: RZ6, preamplifier: RA16PA, differential low impedance amplifier: RA16-LID, TDT Systems) with two foil electrodes positioned in the ear canals (Etymotic) and six cup electrodes (Grass) placed at Fz, Pz, Oz, C7, and both ear lobes, all referenced to ground at the nape. Impedances were kept below 1kΩ by prepping the skin (NuPrep, Weaver and Co.) and applying a layer of conductive gel between the electrode and the skin (EC2, Natus Medical). Stimuli were delivered using calibrated ER3A (Etymotic) insert earphones. Stimulus delivery (sampling rate: 100kHz) and signal acquisition (sampling rate: 25kHz) were coordinated using the TDT system and custom scripts (LabVIEW).

Auditory brainstem responses were measured in response to 3kHz tone pips of 5ms duration. Stimuli had 2.5ms raised cosine ramps, and were presented at 11.1 repetitions per second. Presentation level was fixed at 105dB SPL. Stimulus polarity was alternated across trials and 1000 repetitions per polarity were collected. ABRs from the Fz-tiptrode montage were filtered offline between 300Hz to 3 kHz. Peaks and following troughs of ABR waves were visually marked by an experienced observer, and wave amplitudes were measured using custom written programs (Python).

The FMFR was measured in response to sinusoidal FM stimuli with a carrier frequency of 500 Hz, a modulation rate of 2 Hz and at modulation depths of 0 (i.e. a pure tone), 2, 5, 8, and 10 Hz. The stimuli were 1 second in duration with 5-ms raised cosine ramps and presented once every 1.19 seconds. Stimulus polarity was alternated across trials and 200 samples were acquired for each polarity. The level was fixed at 90 dB SPL. FMFRs from the Fz-tiptrode montage were used for subsequent analyses. Cochlear neural responses to low frequency tones, including the carrier of our FM stimuli, are phase-sensitive in such a way that the summed response to alternating polarities is effectively rectified, and thus periodic at twice the stimulus frequency (Lichtenhan *et al*., 2014). Therefore, we quantified the modulation of the EEG signals with respect to twice the FM carrier frequency, or 1000 Hz. FMFR amplitudes were calculated using a heterodyne method (Guinan *et al*., 2003). Briefly, a discrete Fourier transform (DFT) was computed for each FMFR average. The negative frequency components were discarded to create an analytic signal. This analytic signal was then down-shifted in frequency so that the components around 1000 Hz became centered at 0 Hz. The frequency-shifted signal was filtered in the frequency domain using an exponential filter (Shera and Zweig, 1993), and finally, the inverse DFT was computed. The phase of the inverse DFT is a time-varying signal whose amplitude can be compared directly to the modulation depth of the stimulus. A bootstrapping technique was used to reduce the variability of the calculated FMFR amplitude. An average FMFR was constructed from a subsample of the raw data by drawing 100 samples of each polarity randomly without replacement. This was repeated 1000 times, and the heterodyne analysis was performed on each average. The phase signals output from the heterodyne were averaged and used to compute the final FMFR amplitude. One subject did not yield measurable FMFRs above the noise floor and was excluded from subsequent analyses.

Interaural phase difference following responses were collected to a 520Hz tone carrier whose amplitude was modulated at 40.8Hz. The phase of the carrier was modulated to shift in and out of phase at a rate of 6.8Hz. The degree of shift per ear was 0° (no shift), 22.5°, 45° and 90°. Presentation level was fixed at 85dB SPL. Each stimulus condition was presented continuously for 1.47 minutes, epoched at 294.3ms to contain 300 epochs with two phase shifts each (one out of phase, and one back into phase) and averaged. IPDFR amplitudes and envelope following response amplitudes were calculated from an FFT performed on the averaged waveforms of the Fz-C7 electrode montage at a resolution equal to 1/epoch length (∼3.1Hz), at 6.8 Hz for the IPD response, and at 40.8 Hz for the AM response. Control recordings consisted of phase shifts of 90° in both ears, but in the same polarity to eliminate the binaural component, which showed no responses at the frequency of the interaural phase shift (6.8 Hz).

### Pupillometry

Task-related changes in pupil diameter were collected with a head mounted pupillometry system at a 30Hz sampling rate (Argus Science ET-Mobile), while the subjects used a tablet computer to complete the digits comprehension task (Microsoft Surface Pro 2). The dynamic range in pupil diameter was initially characterized in each subject by presenting alternating bright and dark screens via the tablet computer. Ambient light level was then adjusted to obtain a baseline pupil diameter in the middle of the dynamic range. Pupil measurements were made while subjects were instructed to look at a fixation point on the screen during the presentation of the digits, as confirmed by the experimenter with real time gaze tracking. Pupil data were preprocessed to interpolate for blinks and missing periods using a cubic spline fit, outlier values were removed with a Hampel filter, and the signal was smoothed using a 5-point moving average window. Subjects were excluded if they had extended periods of missing data or if the ambient light could not be adjusted to account for excessive dilation. Reliable data was obtained from 16 subjects who were included for subsequent analyses. A single trial included a 2s pre-trial baseline period, 2s for presentation of the digit sequences, a 2s wait period and the presentation of the virtual keypad to indicate their response. The pupil analysis described here comes from the 2s digit presentation period. Pupil measurements were normalized to the average baseline pupil diameter for each block, collected in the last 500ms of the baseline period. A single measure of pupil-indexed listening effort was operationally defined as the peak value of the average fractional change in pupil diameter function, calculated independently for each SNR block as ((post-stimulus time point – baseline) / baseline). The amplitude value for each SNR was then expressed as a ratio of the complete dynamic range for each subject to reduce variability due to recording conditions, arousal states and eye anatomy.

### Statistical analysis

The distributions of all of the variables were summarized and examined for outliers. Pairwise linear correlations were computed using Pearson’s correlations (r) using custom written scripts in MATLAB. To assess which sets of predictors best account for variability in our outcome measure (Digits comprehension threshold), all predictors were considered in univariable models and the R^2^ calculated (SAS v9.4, SAS Institute). Data from the 16 subjects who had reliable pupil measures were used in the model building, due to the requirement for a balanced dataset across all metrics. Each predictor was then added to the model from highest to lowest R^2^, and the adjusted R^2^ calculated using the formula

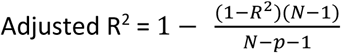

where R^2^=sample R-square, p=number of predictors, N=total sample size. The adjusted R^2^ penalizes for increasing complexity by adding more predictors.

## Acknowledgements

Authors wish to thank William Goedicke and the audiology department of Mass. Eye and Ear for maintaining and providing access to the audiology database. Thanks also to Dr. Jonathan Whitton for help with designing the psychophysical tasks, and Dr. Kelly Jahn for comments on the manuscript.

This study was funded by the National Institutes of Health (NIDCD P50-DC015857) to DBP.

## Author Contributions

AP and DBP conceptualized and designed the study. AP collected and analyzed the data. KEH provided programming support for data collection and analysis. KB and VD provided support for statistical analysis. AP and DBP wrote the manuscript.

---

**Figure 1- Figure supplement 1.**
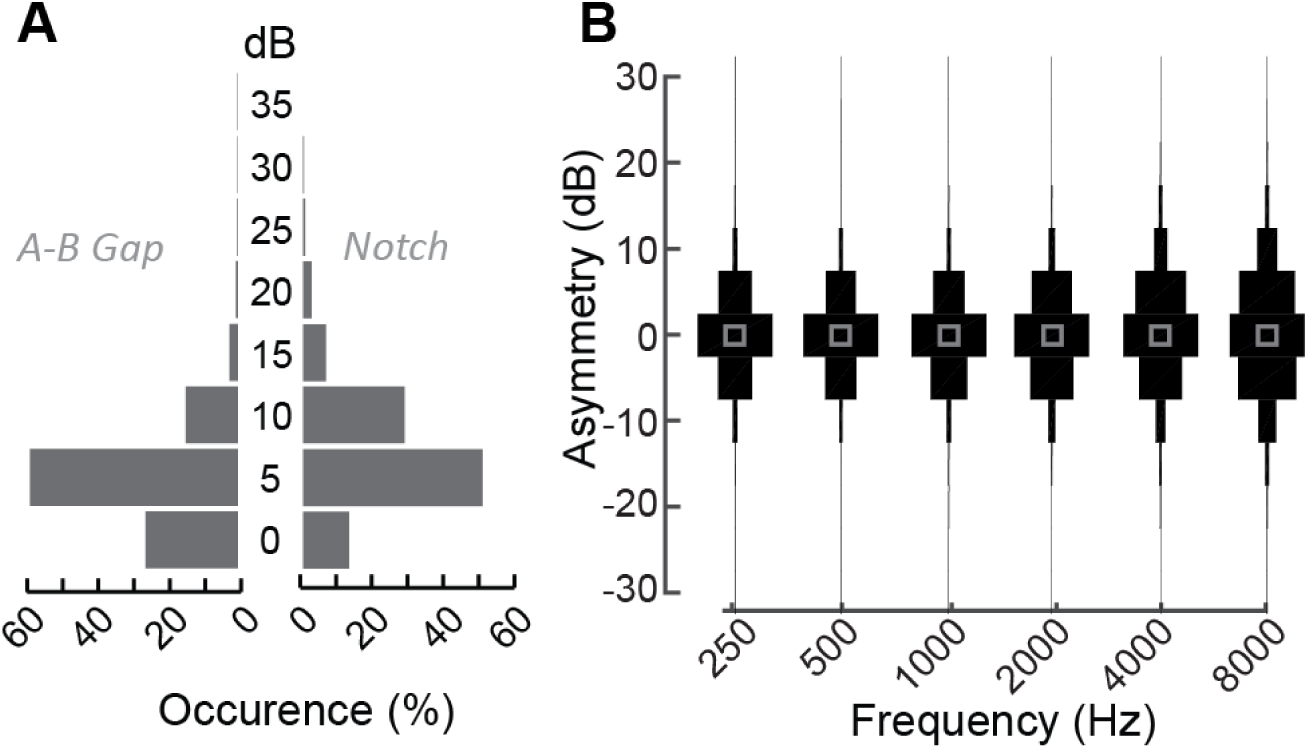
Audiometric characteristics of patients with normal audiograms at the Mass. Eye and Ear audiology clinic. **(A)** Distribution of air-bone gaps, i.e., differences in pure tone thresholds measured via air conduction and bone conduction suggests the lack of any substantive conductive components in these patients with normal audiograms at any test frequency *(left)*. These patients also did not exhibit large focal threshold shifts (notches) in the audiograms, which are indicative of significant noise damage *(right)* (Fausti *et al*., 1981; Mehrparvar *et al*., 2011; Le Prell *et al*., 2013) **(B)** Normalized distribution plots of the difference in hearing thresholds between the right and left ears at the various test frequencies indicate the lack of significant between-ear asymmetries. Hence by all clinical measures of hearing, these patients coming in to the clinic with hearing difficulties are considered to be audiometrically “normal”.

**Figure 1 – Figure supplement 2.**
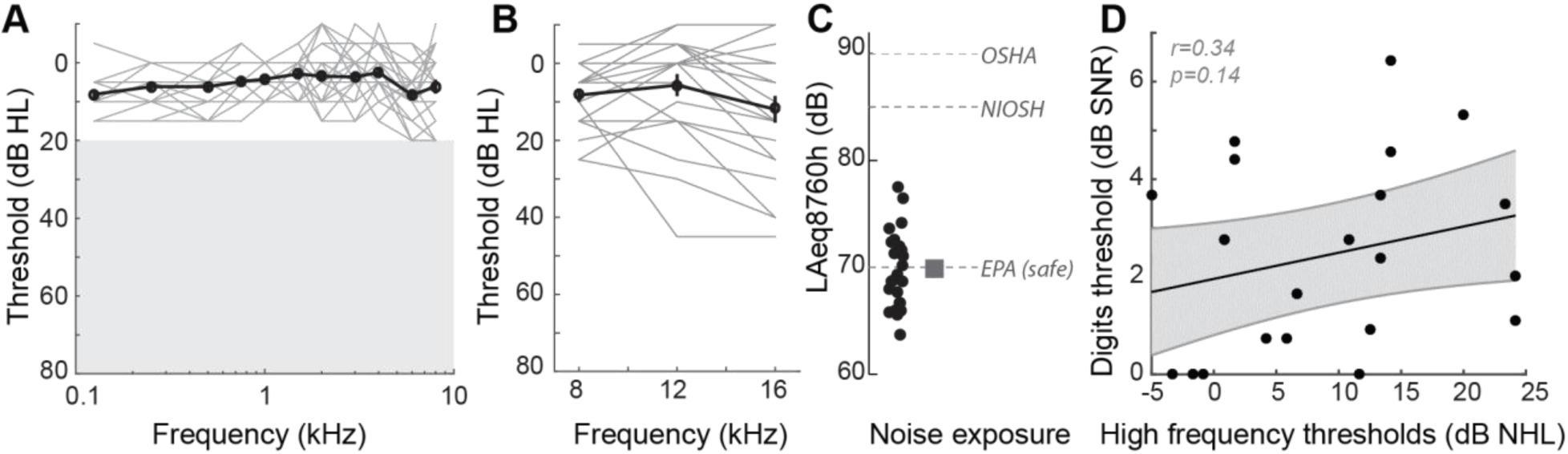
Audiometric profiles and markers of noise exposure in study participants. **(A)** Audiograms from the right ear of the 23 participants in this study, indicating normal hearing thresholds for test frequencies up to 8 kHz. Similar thresholds were present in the left ear (data not shown). Individual audiograms are in gray, and the mean audiogram in black. Unshaded region represents the range of normal thresholds. **(B)** High frequency audiograms, considered an early marker for noise damage showed wide variability in these individuals with normal thresholds in the lower frequencies. **(C)** This came as something of a surprise, as listeners reported lifetime levels of noise exposure that are deemed safe by the EPA, and well below unsafe levels recommended by OSHA and NIOSH. These data suggested that subjective self-reports of noise damage may underestimate the degree of noise damage present in these listeners and that extended high-frequency audibility may be one source of explanation for poor speech processing in noise. **(D)** However, the correlation between the high frequency thresholds and performance on the digits comprehension task was not statistically significant. r = Pearson’s correlation, and shaded area indicates 95% confidence intervals of the regression line (black).

**Figure 1 – Figure supplement 3.**
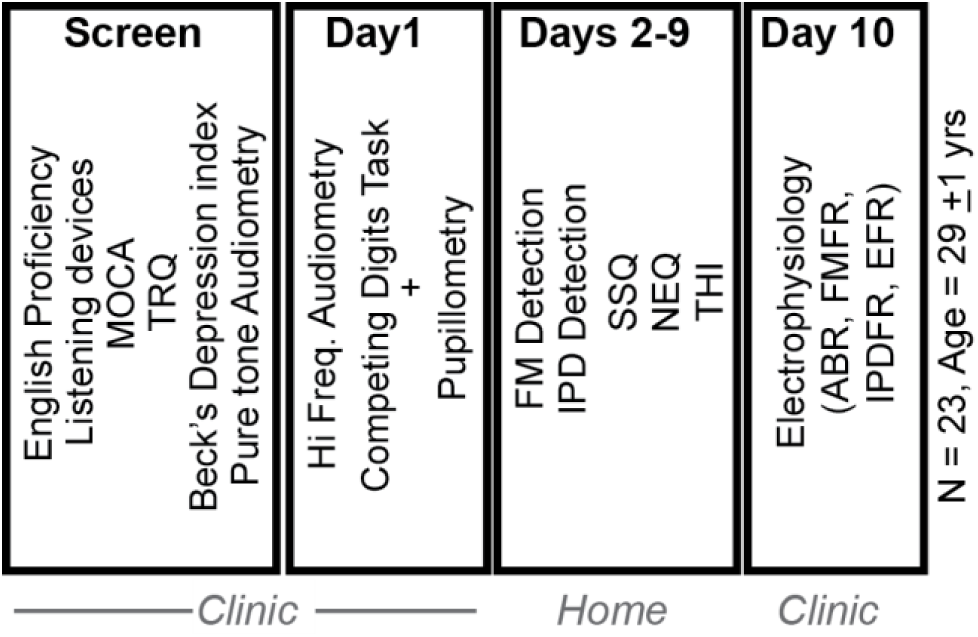
Experimental study design. Of the 27 subjects who provided informed consent to participate in the study, 23 were found to be eligible, based on initial screening for English proficiency, use of listening devices, executive function (Montreal Cognitive Assessment, MOCA), depression (Beck’s depression index), tinnitus (Tinnitus Reaction Questionnaire, TRQ) and pure tone audiometry. Eligible participants completed a set of behavioral and physiological test in the clinic. They were then sent home with tablets that had custom written software, and calibrated head phones to perform additional testing for 8 days. Subjects returned to the clinic with the tablet for a final day of electrophysiological testing.

**Figure 1 – Figure supplement 4.**
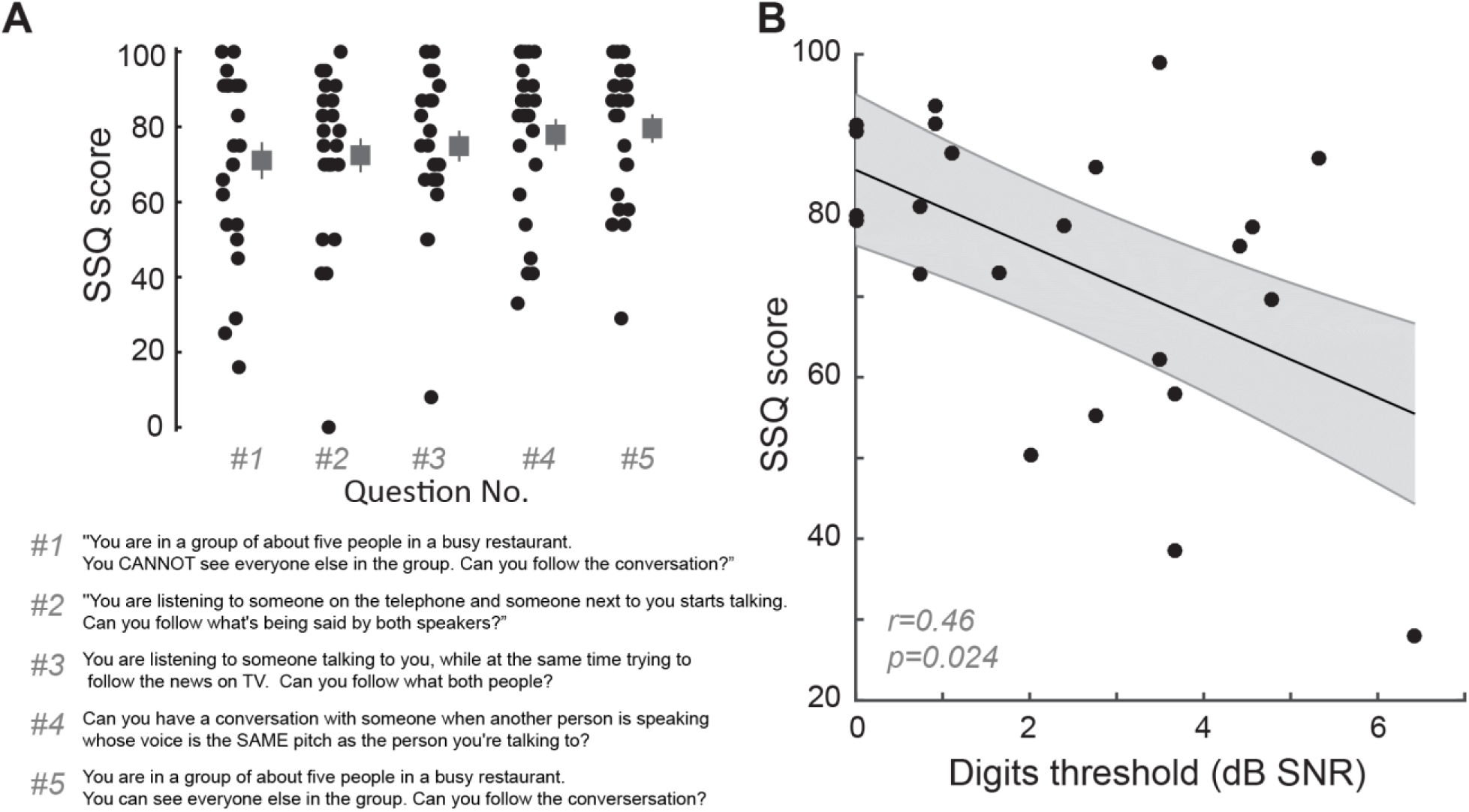
Digits comprehension task captures aspects of self-reported difficulties in real-world multi-talker listening conditions experienced by the participants. **(A)** Questions related to hearing in multiple-speaker situations questions from the speech, spatial and qualities of hearing scale (SSQ) were among the top 5 answers that showed the maximum variability in responses in our participants. Participants answered on a sliding scale with 100 meaning “perfectly” and 0 meaning “not at all”. **(B)** Mean scores on these five questions in the SSQ correlated with the participants’ performance on the digits comprehension task, indicating that the task captures self-reported difficulties of these participants in real world listening scenarios. r = Pearson’s correlation, and shaded area indicates 95% confidence intervals of the regression line (black).

**Figure 2 – Figure supplement 1.**
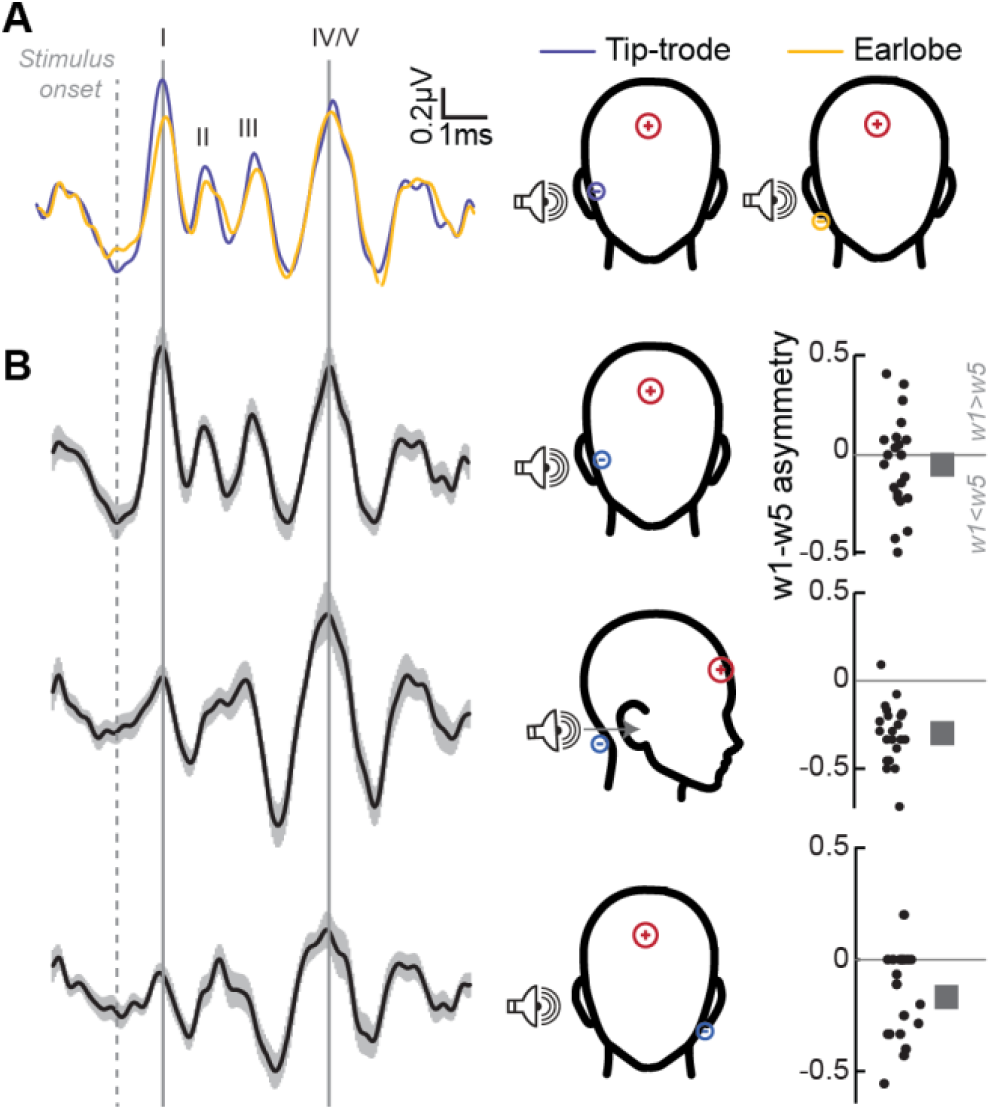
Determination of optimal electrode montages for obtaining electrophysiological responses. **(A)** Grand-averaged ABRs elicited by 3 kHz tones at 105dB SPL, and recorded simultaneously with an electrode on the ear lobe (yellow), and a gold-foil coated ‘tiptrode’ inserted in the ear canal (blue), shows greater amplitudes using tiptrodes for wave 1, with generators in the auditory nerve, but not wave 5, with midbrain generators. **(B)** Simultaneous multi-channel recordings reveal differential contributions from auditory generators for each electrode montage. Grand-averaged ABR waveforms (left) for three electrode montages show the differential contributions of the ABR waves for each montage, reflecting emphasis on peripheral vs. central generators. The relative amplitudes of waves 1 and 5 is characterized by the w1-w5 asymmetry index (right) with values >0 having a larger wave 1, and values <0 having a larger wave 5.

**Figure 4 – Figure supplement 1.**
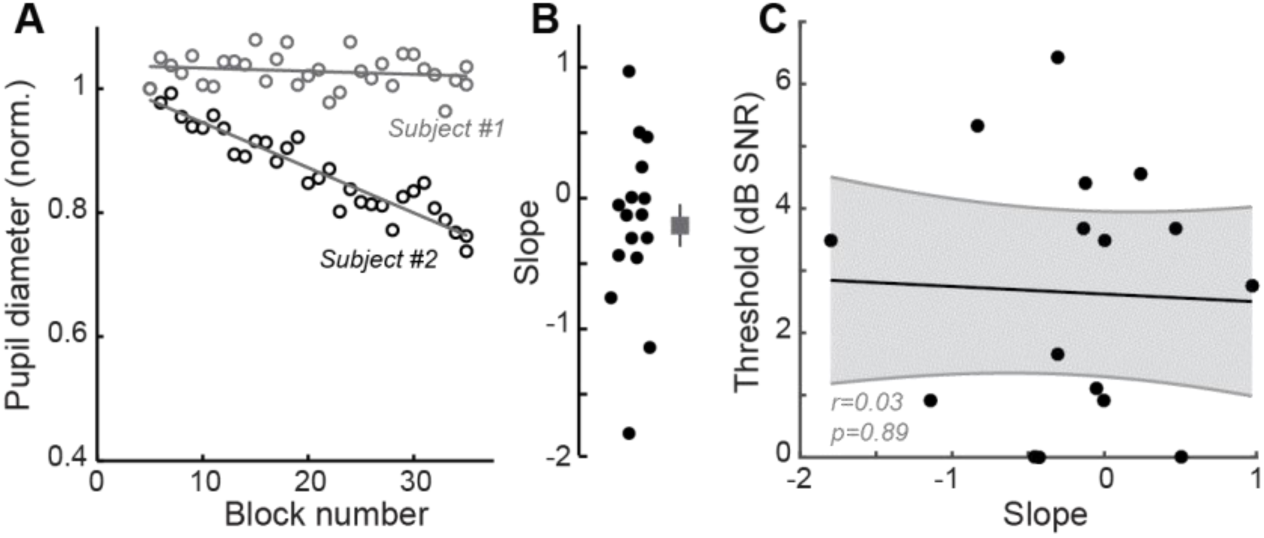
Listening fatigue does not account for performance on the multi-talker digit task. Changes in baseline pupil diameter have been used as an index of listening fatigue (Zekveld, Kramer and Festen, 2010; Zekveld, Koelewijn and Kramer, 2018). **(A)** Baseline pupil diameter measured over the course of successive testing blocks in the 0.5s before trial onset shows minimal changes for one subject and a steady decrease for another. Gray lines show linear fit for each subject. **(B)** Calculated slope of linear fits for all subjects in the study showing the distributions of changes to baseline pupil diameter. Negative values are suggestive of listening fatigue. **(C)** No correlations were observed between changes in baseline pupil diameter and performance on the digits comprehension task, suggesting that listening fatigue did not contribute to changes seen in task performance. r = Pearson’s correlation, and shaded area indicates 95% confidence intervals of the regression line (black).

